# Inferring Cell Differentiation Dynamics with Unobserved Progenitors

**DOI:** 10.64898/2025.12.09.693214

**Authors:** William Howard-Snyder, Richard Zhang, Henri Schmidt, Michelle Chan, Benjamin J. Raphael

**Affiliations:** Dept. of Computer Science, Princeton University, Princeton; 08544 NJ, USA; Lewis-Sigler Institute for Integrative Genomics, Princeton University, Princeton; 08544 NJ, USA; Dept. of Molecular Biology, Princeton University, Princeton; 08544 NJ, USA

## Abstract

A cell differentiation map describes the transitions between cell types during a developmental process, and determining this map is a key challenge in developmental biology. Recent single-cell lineage tracing technologies generate lineage trees that describe the history of cell divisions during a developmental process but do not directly measure differentiation events between cell types. Current approaches to infer cell differentiation maps from cell lineage trees make unrealistic assumptions about the developmental process, do not allow for unobserved progenitor cell types, or do not infer cell-type specific rates of growth and transitions. To address these issues, we introduce TROUPE, a likelihood-based framework that infers differentiation and growth dynamics directly from leaf-labeled cell lineage trees while allowing for unobserved progenitor types via biologically motivated potency constraints. We provide an efficient algorithm for computing the maximum likelihood transition and growth rates, as well as a simple model-selection scheme to choose the number of unobserved types. On simulations, TROUPE recovers transition and growth rates better than previous approaches even when there are unobserved types. We apply TROUPE to a model of mammalian development, Trunk-Like Structures (TLS), and show that TROUPE computes reasonable rates of self-renewal and differentiation in both standard and perturbed conditions.

## 1 Introduction

Understanding the cell differentiation process by which multipotent cells progressively restrict their developmental potential and commit to terminal fates is a central goal in developmental biology. The features of cellular differentiation are often organized as a *differentiation map*: a graph where the nodes represent cell types and edges denote permitted transitions between them. But missing from this definition of the cell differentiation map is the notion of *rates* at which cell types differentiate and grow, which is crucial to the dynamics of the process. Although the cell differentiation maps and their rates cannot be directly measured, recent lineage tracing technologies [1–10], when paired with tree reconstruction methods [11–15] enable the construction of high-resolution, high-throughput cell lineage trees that describe the history of cell divisions during development in complex organisms. Further, shared heritable markers can be used time-resolve these trees, providing times between cell divisions. These time-resolved lineages with transcriptionally-annotated leaves provide the opportunity to infer cell differentiation maps and their rates.

Current methods for inferring cell differentiation maps from lineage trees do not rigorously infer growth and transition rates. Instead, methods cluster cell types based on pair-wise distances defined over cell lineage trees [7, 16–18], use cell lineage trees to inform cellular trajectories [19], or optimize parsimony-like criteria [20]. Some of these methods [16, 20] give transition counts between cell types which offer a sense of the frequency of differentiation between cell types, but these counts are confounded by the simultaneous effects of differentiation and division, and thus are not cell-type specific rates of growth and differentiation. However, as we show below, we can infer these rates from cell lineage trees that have both time-resolved branch lengths and cell type annotations at their leaves. One recent method attempts to infer rates for developmental processes [21], but this method cannot infer unobserved progenitor types and also makes unrealistic assumptions about the developmental process such as that all cell type transitions are allowed.

To infer the differentiation dynamics from cell lineage trees, one needs an explicit model of cell growth and transition on these trees. The State Speciation and Extinction (SSE) framework – including BiSSE [22], MuSSE [23], HiSSE [24], QuaSSE [25], as well as adaptations to handle incomplete sampling of extant taxa [26–29] – models how traits (e.g., cell types) evolve on a tree over time. These models allow for the inference of growth and transition rates of types over the tree given types observed at the leaves via maximum likelihood estimation, and thus are a good fit for the data available from cell lineage trees [30]. However, existing SSE models are missing two key features of development: (1) irreversible transitions that restrict in differentiation potential and (2) progenitor cell types that may not be observed among the extant taxa.

We introduce the Potency-constrained State Speciation and Extinction (PoSSE) model, which models cell differentiation and growth in a way that respects the rules by which cell types transition during development. Specifically, PoSSE reflects the gradual restriction of *potency* over time. Early in development, cells and their progeny can differentiate into many cell types, but over time, they commit to increasingly restricted sets of cell fates. We formally encode potency as a subset of the terminal cell types and require that cell types of one potency transition only into cell types whose potency is a subset of their own [20]. Using PoSSE we derive sufficient conditions under which certain progenitor cell types have vanishing probability of being observed at sampling time. These unobserved cell types pose challenges for rate inference, and complicate the identification of the number of progenitors. Leveraging potency constraints decreases the number of parameters in the process, improving our ability to infer rates when intermediate progenitors are unobserved. Potency constraints also clarify the role of such unobserved types: they can be transiently occupied types that influence growth and composition while not being present at the time of observation.

We also introduce *Tree-informed Rate Optimization Using Potency Enforcement* (TROUPE), an algorithm for computing the maximum likelihood estimator of PoSSE parameters from leaf-labeled trees. To address unobserved progenitors, TROUPE includes a simple model-selection procedure that chooses the number of unobserved progenitor cell types. The algorithm scales to large collections of trees with many leaves and jointly infers rates of cell differentiation and growth. On simulations, TROUPE recovers transition and growth parameters more accurately than existing approaches, including in settings with unobserved progenitors. We apply TROUPE to Trunk-Like Structures (TLS), a stem-cell–derived *in vitro* model of early mammalian trunk development, grown under standard and perturbed conditions. The inferred rates of differentiation and growth recapitulate the expected model of development and reveal subtle, but consistent changes in differentiation rates between conditions. By directly modeling cell growth and differentiation, we obtain interpretable estimates of cell-type-specific transition and growth rates as well as expected population composition over time. These insights facilitate a deeper understanding cell population dynamics in developing organisms.

## 2 A Branching Process Model of Development

We model cell proliferation and differentiation in a given biological system as a continuous-time, time-homogeneous, *n*-type, pure-birth branching process, i.e. a multi-type Yule process. Each cell has a latent type in Ω, the set of possible cell types in this system. This branching model is parameterized by a transition rate matrix ***Q*** ∈ ℝ^*n×n*^ where entry ***Q***_*i,j*_ governs the rate at which type *i* transitions into type *j*; growth rate parameters **λ** ∈ ℝ^*n*^, where entry **λ**_*i*_ governs the rate at which type *i* gives birth to a new cell of the same type; and an initial distribution ***π*** where ***π***_*i*_ denotes the probability that the process starts from a cell of type *i* [31, 32]. This class of models is widely used for studying cell populations (e.g., hematopoiesis, cancer, cell-cycle) [33–35], but – to our knowledge – has not been applied to model developmental systems with many types and terminal fates (e.g., embryogenesis).

In developmental systems, cell types exist in a hierarchy of increasingly restricted potential or *potency*; early progenitor cell types can differentiate into many fates whereas later types with restricted potential can access only a subset. Additionally, cells do not often revert to previous types with higher potency. The dynamics of type transitions in this system are naturally *reducible* as there exist absorbing, fully differentiated terminal types. In this section we define the PoSSE model, which encodes the hierarchy of cell types in development into the framework of multi-type branching models via constraints on the rate matrix ***Q***. We further show that a consequence of the reducible dynamics of this system is that intermediate progenitors may be rare or absent among sampled leaves at realistic observation times, which complicates inference and, if ignored, can bias rate estimates.

### 2.1 Tree generative process

For fixed parameters (***Q, λ, π***), the branching process induces, at any observation time *t >* 0, a random rooted binary ultrametric tree *T* = (*V, E, b*) (vertices, edges, and branch lengths) with latent cell types ***y*** = (***y***_*v*_)_*v*∈*V*_. The process starts from a single root cell at time 0 whose type ***y***_*ρ*_ is drawn from the initial distribution ***π***. Each cell of type *i* divides at rate *λ*_*i*_, producing two daughters of the same type and creating a binary split in the tree. The length of each edge corresponds to the time until division. Between division events, the type of each cell evolves as a continuous-time Markov chain on Ω with generator ***Q***. Equivalently, along each edge *e* ∈ *E* of length *b*_*e*_, the type trajectory can be viewed as a coloring *c*_*e*_ : [0, *b*_*e*_] → Ω given by this Markov chain started at the parent’s type. Conditioned on their current types, different lineages evolve independently.

At time *t*, when the process is observed (e.g., the end of a lineage tracing experiment), we obtain a fully labeled ultrametric tree *T* whose leaves correspond to the cells alive at time *t* with types ***y*** = (***y***_*v*_)_*v*∈*V*_. In a lineage-tracing experiment we observe the tree topology and branch lengths together with the types of the leaves only, so the data consist of leaf labels ***x***_*v*_ = ***y***_*v*_ for each leaf *v*, while internal node types and within-edge colorings {*c*_*e*_} _*e*∈*E*_ are unobserved. The empirical type composition of the leaves is therefore identical in distribution to the population type composition of the underlying branching process at time *t*. We provide an explicit construction of the tree in Section S7, which is similar to the construction in Dragomir et al. [36].

### 2.2 Likelihood

Given a rooted tree *T* = (*V, E, b*) with *m* leaves labeled ***x*** ∈ Ω^*m*^ for some size *n* label-set Ω. Under the tree generative process described in the previous section, the likelihood of *T* given the transition rates ***Q*** ∈ ℝ^*n×n*^, the growth rates ***λ*** ∈ ℝ^*n*^, and the root distribution ***π*** is:

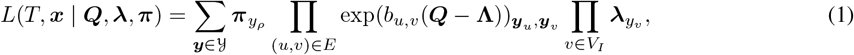

where *ρ* indexes the root node, *V*_*I*_ are the internal nodes of *V* (i.e., nodes that are not leaves), 𝓎 is the set of labelings of *V* that extend ***x***, and **Ω** = diag(***λ***). This likelihood is identical to that of the BiSSE [22] or MuSSE [23] models with the death rate set to zero. While there are reports of cell death in organismal development, the death rate is substantially smaller than growth and setting death rate to zero is a reasonable approximation. We emphasize that the death rate does not need to be set to zero in this model, although non-zero death rates do not have a closed form expression for the likelihood making optimization substantially more challenging.

For a set 𝒯 = {(*T* ^(1)^, ***x***^(1)^),…, (*T* ^(*N*)^, ***x***^(*N*)^)} of *N* independent leaf-labeled trees from the same developmental system (***Q, λ***). The likelihood of τ is

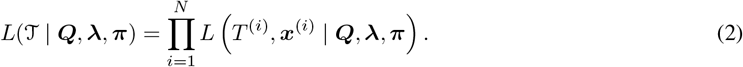

### 2.3 Potency-constrained State Speciation and Extinction (PoSSE) Model

The *potency* of a cell type is the set of all terminal cell types that it or its progeny can differentiate into. We give a formal description of the notion of potency by first identifying a subset *S* of the cell types that are terminal in a given developmental process, i.e. they cannot differentiate any further. The potency *α* of a cell type is a subset of *S*, i.e. an element of the power set 𝒫 (*S*) of *S*. We require that the potency of a cell uniquely identifies its type, so that each cell type is defined by its potency. To reflect the notion that cells increasingly restrict their potency over time, cell type *α* ∈ 𝒫 (*S*) can transition into a cell-type *β* ∈ 𝒫 (*S*) only if *α* ⊇ *β*. This is similar to the notion of potency in Sashittal et al. [20], however, we include only terminal types, rather than all observed types. Since cell types are defined by their potencies, the transition rate matrix ***Q*** describes transitions between elements of *P* ⊇ 𝒫 (*S*) (taking Ω to be *P*), where *P* a set of potencies. The following definition translates the constraint that potencies can only transition to subsets of themselves into constraints on the parameters ***Q***.

#### Definition 2.1.

*The rate matrix* ***Q*** ∈ ℝ^*n×n*^ *is* potency-constrained *with respect to the set of potencies P ⊇* 𝒫 (*S*) *if for any two potencies α, β ∈ P, if α* ⊉ *β, then Q* _*α, β*_ = 0.

The PoSSE model combines these potency constraints on ***Q*** with the likelihood defined in Equation 1. Namely, for potency-constrained transition rate matrix ***Q*** the likelihood of ***Q, λ, π*** under PoSSE is given by Equation (1) and if ***Q*** is not potency-constrained then the likelihood is 0. We further assume that the starting state has the largest potency. In terms of our parameters, this means that ***π***_*S*_ = 1 and for all *α* ∈ *P, *α* ≠ S*, we have that ***π***_*α*_ = 0. Note that under PoSSE, then any tree labeling that violates the potency constraints (i.e., descendant potency is not a subset of ancestor potency) will have a likelihood of 0.

For any potency *α* ∈ *P* that corresponds to a terminal state, we have | *α* | =1 because the terminal states are the states that cannot differentiate further (i.e., are only potent for themselves). If ***Q*** is potency-constrained with respect to *P* and *P* contains *k* terminal potencies, the rows corresponding to those terminal potencies will be all zeros. Thus, we can write it as

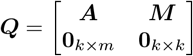

where **0**_*k×m*_ is the all zeros matrix with *k* rows and *m* columns. We use the convention that larger potency states index the earlier rows of ***Q*** (e.g., the first row gives the outgoing transitions of the most potent type). So, ***Q*** will be an upper triangular matrix. When treated as a weighted adjacency matrix, the resulting graph forms a DAG, which can be interpreted as the differentiation map for the developmental system where edges are weighted by their transition rates (Figure 1).

**Fig. 1:**
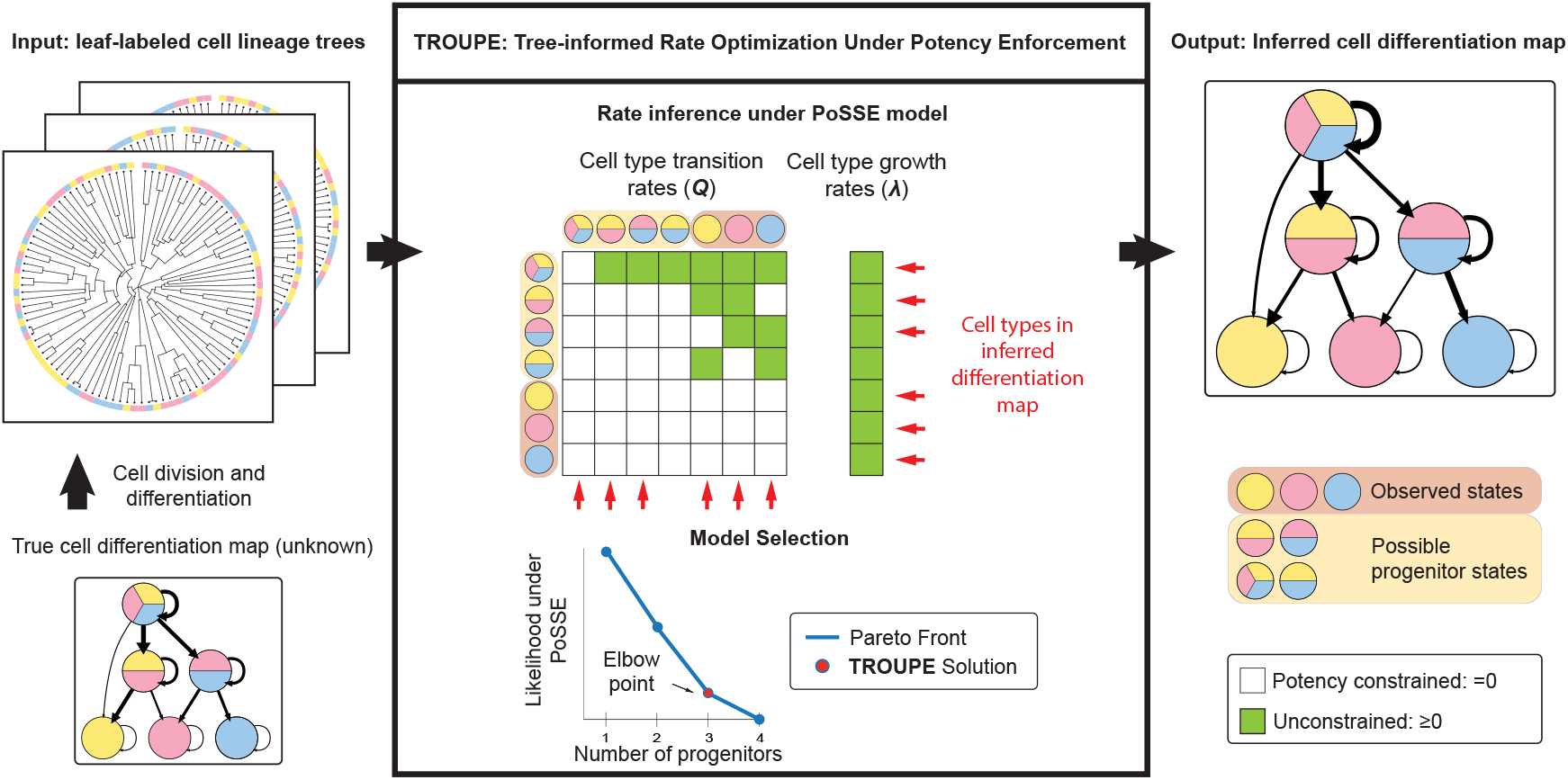
Overview of TROUPE. TROUPE takes as input a set of cell lineage trees whose leaves are labeled by cell types. These trees are assumed to have been generated according to some true cell differentiation map under the PoSSE model with true growth and transition rates. TROUPE then infers the rates of transition between the terminal and the possible unobserved progenitors as defined by their potency. The PoSSE model imposes constraints so that cell types can only transition to ones with more restricted potency. TROUPE then performs model selection by generating the Pareto front showing the tradeoff between the likelihood and the number of progenitors in the differentiation map. The differentiation map corresponding to the elbow of the Pareto front is returned.

### 2.4 Unobserved Cell-Types

Progenitor types may be unobserved in branching processes with potency-constrained transition matrices. We can see how this can occur by analyzing the expected population composition over time. Let **Ω** = diag(***λ***) ℝ^*n×n*^. The (*i, j*) entry of the matrix exponential below gives the expected number of types in type *j* after time *t* if the process starts with a single cell of type *i* [31, 32]

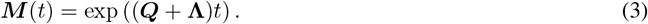

Suppose there are *n* =2 states, with state 1 being a progenitor and state 2 being a terminal state. Additionally, suppose that the process starts with *r* cells of the most potent type. Then, the number of progenitors decays exponentially in time under certain conditions.

#### Proposition 2.1.

*Let *α* be the outgoing transition rate of the progenitor type and ω*_1_ *be its growth rate. If *α* > λ*_1_, *then the expected number of progenitor types over time is r****M*** (*t*)_11_ *< r* exp (*t*(*λ*_1_ *− α*)).

*Proof*. See Section S8.1 □

The general case of *n >* 2 cell-types reduces to the 2-state case by treating the totipotent cell-type as the progenitor state and all other types as terminal states. This gives us the immediate corollary regarding the expected decay of the totipotent type in an *n*-type potency-constrained branching process by 2.1.

#### Corollary 2.1.

*Let i index the totipotent type and α*_*i*_ = *Σ*_*j≠i*_ ***Q***_*ij*_ *be its outgoing transition rate. If α*_*i*_ *> λ*_*i*_, *then the expected number of totipotent types over time is r* exp (*t*(*λ*_*i*_ *− α*_*i*_)).

For the general case with *n >* 2 states, the expected number of *all* progenitors vanishes in the long time limit if the outgoing transition rate for each progenitor *i* exceeds its growth rate i.e., ***λ***_*i*_ *< Σ* _*j≠ i*_ ***Q***_*ij*_ = *α*_*i*_. That is,

#### Proposition 2.2.

*Let z*(*t*) *be the number of progenitor types at time t (i.e., types i such that* ***Q***_*ii*_ *< 0). If α*_*i*_ *>* ***λ***_*i*_ *for each progenitor type i, then* 𝔼 [*z*(*t*)] *→* 0 *as t → ∞*.

*Proof*. See section S8.2.□

In the following section, we describe our method for identifying the number of unobserved progenitors and inferring their transition and growth rate parameters.

## 3 Maximum Likelihood Estimation under PoSSE model

In this section, we introduce TROUPE, an algorithm to perform maximum likelihood inference under the PoSSE model. The potency constraints makes computing the maximum likelihood estimator of (***Q, λ***) under the PoSSE model a constrained optimization problem that is more challenging than under the unconstrained SSE models. We derive TROUPE through a series of intermediate optimization problems.

We are given as input a set 𝒯 = {(*T* ^(1)^, ***x***^(1)^),…, (*T* ^(*N*)^, ***x***^(*N*)^)} of *N* leaf-labeled trees from the same developmental system. Since ***Q*** and ***λ*** are intended to be *valid* transition and growth rates, assume the following constraints are present in every optimization problem: for each row *i* we have ***Q***_*ii*_ = *− Σ* _*j ≠i*_ ***Q***_*ij*_, ***Q***_*ij*_ *≥* 0 for *j ≠ i*, and ***λ***_*i*_ *≥* 0.

### 3.1 Observed progenitors (SSE model)

Suppose that *all n types have been observed among the leaves*. Then, inference of the transition and growth rates by maximum likelihood under the SSE model can be accomplished by solving

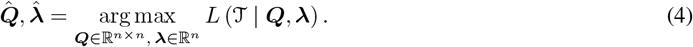

Since the loss is differentiable and the feasible set can be re-parameterized so that it is unbounded (e.g., with softplus parameterization of the off-diagonal entries of ***Q***) and we can employ gradient-based optimization solvers to solve this optimization problem. In particular, we use L-BFGS [37].

Solving for the maximum likelihood estimator under the standard SSE is insufficient for development for two reasons: (1) not all cell types are observed as progenitor cell types disappear under the reducible dynamics of development and (2) potency restrictions are not enforced in how cell types can transition to each other. These factors complicate rate inference, and we explore this difficulty empirically in Section 4.1. To better infer the parameters in this setting, we describe an optimization problem that enforces the potency constraints described in Section 2.3.

### 3.2 Fixed-number of unobserved progenitors

To build intuition for our final procedure, we start with an optimization problem that requires an assumption that the true number of cell types is known, but enforces potency constraints and infers the potencies of unobserved cell types. Suppose that the total number of cell types is *n*, of which only *m ≤ n* have been observed. Additionally, suppose we know the potency of each observed type. Let *B ⊆* 𝒫 (*S*) be the set of observed potencies. Note that for each observed terminal type *x* ∈ *S* we have that {*x*} ∈ *B*. We seek the maximum likelihood estimates 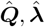 such that ***Q*** is potency constrained with respect to a size-*n* potency set *P*,

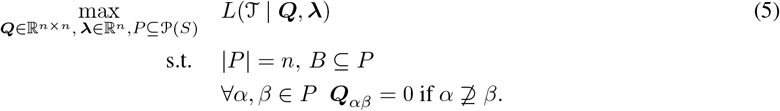

This constraint ensures that ***Q*** is potency constrained with respect to a set of potencies *P*.

Further restricting to rate matrices that are potency-constrained, the feasible set is non-convex and we cannot solve the optimization problem directly with gradient-based solvers. The brute-force solution would involve solving a separate MLE for each of the size *n* potency sets *P* that contain the observed potencies *B*. There are 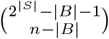 such potency sets (2^|*B*|^ *−* |*B*| *−* 1 is the size of the set 𝒫 (*S*) \ (*B ∪ ∅*)). For |*S*| = 4 or less terminal states, this can be solved relatively quickly, but for 5 or more terminal states it becomes intractable (e.g., with *n* =9 types, |*S*| =5 terminals, and *B* = *S* observed progenitors, we have to solve *≥* 10^9^ independent MLE problems).

### 3.3 Potency constrained MLE as column-selection

To address the scalability issues of the optimization problem in Equation (5), we start by recasting the problem as a column-selection problem inside an over-parameterized model. Rather than optimize over all potency sets, we can fix a large potency set *P′*, constrain ***Q*** with respect to it, and require that at most *n* columns be non-zero. Specifically, let *P′* = 𝒫 (*S*) \ ∅ to be the largest potency set and let *q* = |*P*|. Let ***Q***_*i*_ be the *i*th column of ***Q***, and let ‖***Q***_*i*_ ‖_0_ = 0 if ***Q***_*i*_ = **0** and 1 otherwise.

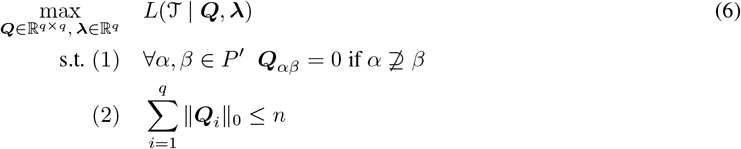

#### Proposition 3.1.

*Equation* (5) *is equivalent to* *Equation* (6).

*Proof*. See section S8.3. □

Constraint (2) makes Equation (6) difficult to solve. The feasible set is still non-convex and enumerating over all possible ways to have at most *n* non-zero columns is just as intractable as enumerating over all possible potency sets. Rather than impose constraint (2) explicitly, we use *L*1-regularization to encourage the columns to be sparse. That is, we fix a regularization strength γ*>* 0 and solve

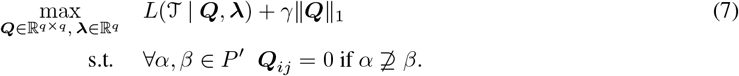

The potency set is fixed and not treated as a parameter to be optimized (unlike Equation (5)), which allows us to parameterize ***Q*** so that the feasible set is unconstrained and apply gradient-based solvers.

This formulation trades likelihood for model complexity through the regularization strength parameter *γ*. Let 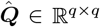 be the solution to Equation (7), where *q* is typically much larger than desired. For sufficiently large values of *γ*, many types will be unreachable from the totipotent state in the differentiation map induced by 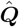. Let the smaller rate matrix 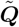 consist of the rows and columns of 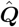 that correspond to just these reachable types. Similarly, let 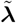 be only the entries of 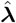 that correspond to reachable types in 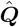. The reachable-type matrix 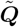 is potency-constrained according to the potencies 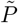 of the reachable types, however, the rates will be biased due to regularization. To correct for this bias, we fix the potency-constraints inferred by 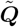 and optimize under just the likelihood criterion (Equation 5) without any regularization fixing 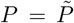. These steps are summarized in steps (i)-(ii) in the inner for loop of Algorithm 1 in Section S9

Note that this does not guarantee that our rate matrix is *n ×n* for some pre-specified *n*. In practice, we solve Equation (7) using several different regularization strengths γ, and find that for sufficiently fine-grained values of *γ*, there is often a 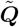 that realizes *n* types for desirable values of *n*. In the next section, we describe our approach for identifying *n*.

### 3.4 Finding an unknown number of unobserved progenitors through TROUPE

The overparameterized formula in Equation (7) allows for inference under potency enforcement in a scalable way, but does not select the number of unobserved progenitors. Here, we develop an algorithm which involves solving the previous optimization problem as a subroutine to determine the number of unobserved progenitors.

Let Γ = {*γ*_1_ *> γ*_2_ *>* … *> γ*_*K*_} be a decreasing sequence of regularization strengths (we find that log-spacing the values works well in practice). For each *γ* ∈ *Γ*, we solve Equation 7 and debias to obtain transition and growth rate estimates 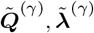. We also record the model size *n*^(*γ*)^ = |*P* ^(*γ*)^ | and fit log *L*(𝒯 | ***Q***^(*γ*)^, ***λ***^(*γ*)^).

Since models with a larger number of types *n*^(*γ*)^ can match or exceed the likelihood of ones with fewer types, raw likelihood comparison is uninformative. Therefore, we form the following set of pairs:

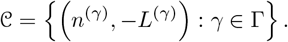

We then compute the Pareto front of 𝒞 i.e., points for which no model achieves both a smaller number of types *n* and a lower negative log-likelihood.

On the Pareto front, we select the elbow where marginal likelihood gain per additional type begins to diminish. Concretely, we sort the Pareto points by increasing *n*, apply the Kneedle [38] procedure to find the index of maximum deviation from the line joining the endpoints (i.e., maximum curvature for discrete data), and take the corresponding regularization strength *γ*^⋆^. These steps are summarized in Algorithm 1 of Section S9. This is analogous to the Carta [20] selection criteria, but applied to likelihood rather than discrepancy.

The procedure returns the final debiased estimates 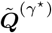 and 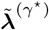. This selection layer completes TROUPE: regularized over-parameterization to explore candidate potency sets, debiased refitting to obtain unbiased likelihoods, and a simple reproducible rule to pick the number of unobserved progenitors.

## 4 Results

### 4.1 Evaluation on simulated data

We compare TROUPE, SSE, *Carta*, and Evolutionary Coupling (EvoC) in terms of each method’s ability to recover the ground-truth cell differentiation map on simulated data. *Carta* is a method that does take into account potency conditions in inferring cell differentiation maps and finds the most parsimonious set of potencies (progenitor cell types) for observed cell lineage trees. Evolutionary Coupling represents a class of methods that infer cell differentiation maps by clustering cell types based on pair-wise distances on the cell lineage tree. Methods of this class are only able to produce a much restricted subset of the possible cell differentiation maps that can be produced on a set of terminal states, specifically binary tree cell differentiation maps in which the number of progenitors is |*S*| *−* 1.

To generate our simulated instances, we define a ground-truth cell differentiation map with fixed parameters (***Q***^⋆^, ***λ***^⋆^). We visualize (***Q***^⋆^, ***λ***^⋆^) as a weighted graph (Fig. 2a). We simulate *N* = 16, 32, 64, 128, 256 independent cell lineage trees from a branching process that proceeds according to (***Q***^⋆^, ***λ***^⋆^), which we stop after a fixed cutoff time of 2.35. For each *N*, we repeat this procedure 5 times to get 5 independent trials, and for each trial we take the simulated cell lineage trees as input to each method. We then compare the inferred cell differentiation map of each method for each trial to the ground-truth cell differentiation map according to two criteria: 1) the ability to recover the ground truth set of progenitors and 2) the rates of transition and growth for those methods that infer rates (TROUPE and SSE). For each method, we provide the number of ground-truth progenitors (4) as additional information as only *Carta* and TROUPE have defined procedures to determine the number of ground-truth progenitors.

**Fig. 2:**
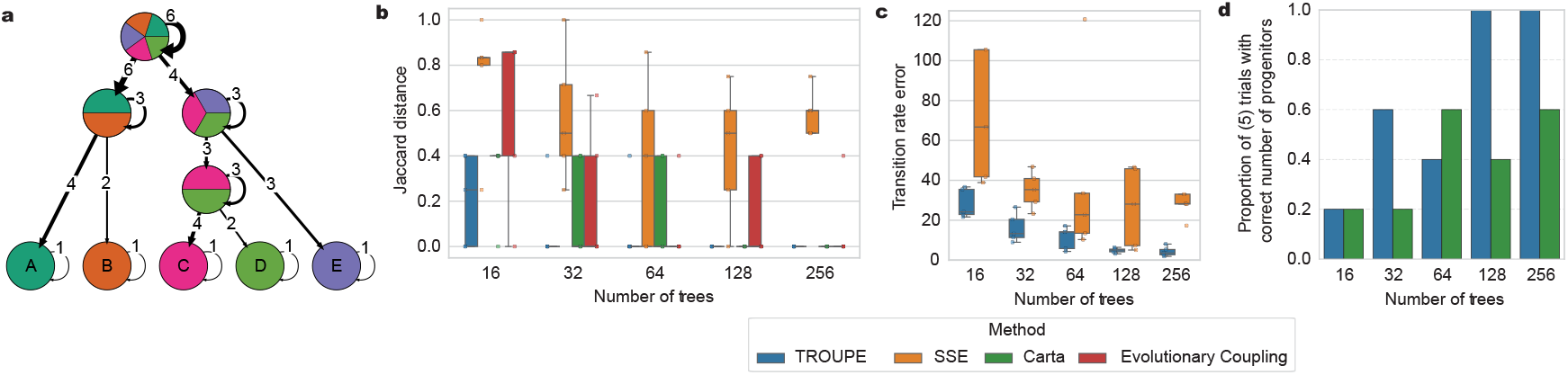
TROUPE outperforms other methods in inferring the ground-truth progenitors and rates in simulation. (a) The ground-truth differentiation map used in these simulations. Each node represents a row of ***Q***^⋆^, the edge weight from node *u ≠ v* is ***Q***_*u,v*_, and the self-edge weight for node *v* is given by ***λ***_*v*_. (b) Jaccard distance between the set of ground-truth progenitors and each set of inferred progenitors by each method for 5 trials for each number of trees. (c) Absolute error in transition parameter values for SSE and TROUPE for 5 trials for each number of trees. (d) The proportion of 5 trials for which the model selection procedure of TROUPE and *Carta* determine the correct number of progenitors in the ground-truth progenitor set.

TROUPE better recovers the ground-truth set of progenitors in these simulations than other methods. TROUPE, EvoC, and *Carta* achieve Jaccard distance of 0 for large *N* (256) meaning they perfectly infer the ground-truth progenitors across all trials (Fig. 2b). SSE does not achieve this accuracy, even for large *N*. TROUPE achieves 0 Jaccard distance with many fewer cell lineage trees (32) as compared to EvoC (64) and *Carta* (128), suggesting superior sampling efficiency of TROUPE. Although EvoC achieves 0 distance across all trials for *N* = 64, it does not for *N* = 128. For *N* = 16, 32 trees, TROUPE has lower mean of Jaccard distance (16: 0.21, 32: 0.08) than *Carta* (16: 0.32, 32: 0.16) and EvoC (16: 0.59, 32: 0.21), indicating that it is able to better infer the correct progenitors for these numbers of trees (Fig. 2b).

For each number of trees, the parameters inferred by TROUPE (both the growth and transition rates) are closer to the ground-truth parameters than SSE. To measure this, we compute the total difference in transition and growth rates between those inferred by TROUPE (or SSE) and the ground-truth rates. For *N* = 256, TROUPE achieves 4.3 mean transition rate error and 1.2 mean growth rate error compared to 58.0 mean transition rate error and 6.51 mean growth rate error for SSE (Fig. 2c, Fig.S1). For both methods, the absolute error tends to decrease as the number of trees increases. However, SSE requires many more trees to achieve the same level of transition rate error as TROUPE. For example, at 32 trees we achieve an average transition rate error of 16.0 which is lower than the SSE average transition rate error for all *N* (Fig. 2c).

We also evaluate how well *Carta* and TROUPE identify the correct number of progenitors in the underlying ground-truth cell differentiation map. We use the same set of simulated cell lineage trees. These simulations confirm the above result that branching processes that follow potency-constrained transition matrices will have progenitor cell types that will go unobserved, with a mean of 0, 0, 0.4, 0.4, 1.6, 1.8 progenitors being observed at the leaves (out of the 4 ground-truth progenitors) over 16, 32, 64, 128, 256, 512 trees, respectively. Hence, the true number of progenitor cell types need to be inferred as we do not observe all progenitors even for large numbers of trees.

For TROUPE, the number of progenitors is inferred as described above in section 3.4 over regularization strength *γ* ∈ {0, 0.001, 0.003, 0.01, 0.03, 0.1, 0.3, 1, 3, 10} and for *Carta* we infer the knee from *k* = 3, 4…, 7 unobserved progenitors. TROUPE is better able to infer the true number of progenitors in the ground-truth cell differentiation map than *Carta*. Over 5 trials for each of *N* = 128, 256, TROUPE is able to infer the proportion with perfect accuracy (Fig. 2d). For all *N, Carta* is unable to achieve greater than 0.6 accuracy of 5 trials. While *Carta* claims that its proposed model selection framework infers the true number of progenitors, its accuracy is low. TROUPE shows that this model selection framework is viable and can infer the true number of progenitors when the Pareto front is calculated with a likelihood objective.

### 4.2 TROUPE infers differentiation dynamics in a model of mammalian development

We apply TROUPE to cell lineage trees inferred from CRISPR-Cas9 lineage tracing of the mouse Trunk-like Structure (TLS) gastruloid model [18, 39], an *in-vitro* model of mammalian gastrulation as well as a perturbed condition of this model. In TLS, individual structures are grown from embryonic stem cell aggregates that are induced to differentiate and form trunk structures that resemble precursors to the spinal cord and surrounding muscle-like tissues. In the elongating tip of each structure, a pool of self-renewing neuromesodermal progenitors (NMPs) continuously differentiate into somite and neural tube cells (Fig. 3a). In a perturbation condition of TLS named TLS^CL^, two small molecules – a WNT agonist (**C** for CHIR99021) and a BMP inhibitor (**L** for LDN-193189) – are applied during TLS formation. This perturbation leads to a clear morphological phenotype that shows excess somitic production (Fig. 3a), but the exact changes to the rates of differentiation and growth of somites relative to neural tube cells are unknown. These change in rates are key to understand the degree of this effect. In addition to the NMP cells and their descendants, other primitive streak derivative cell types – such as endothelial, endoderm, and primordial germ-cell-like cells (PGCLCs) – are also observed in this system (Fig. 3b). PGCLCs, which are derived from the epiblast and are not part of the three major somatic germ layers, commit first. The origin of and differentiation dynamics of the endothelial and endoderm cells are unclear after the germ layer, with one putative model being a separation of the mulitpotent pool of the germ layer into progenitors that are biased in the formation of these respective cell types (Fig. 3b).

**Fig. 3:**
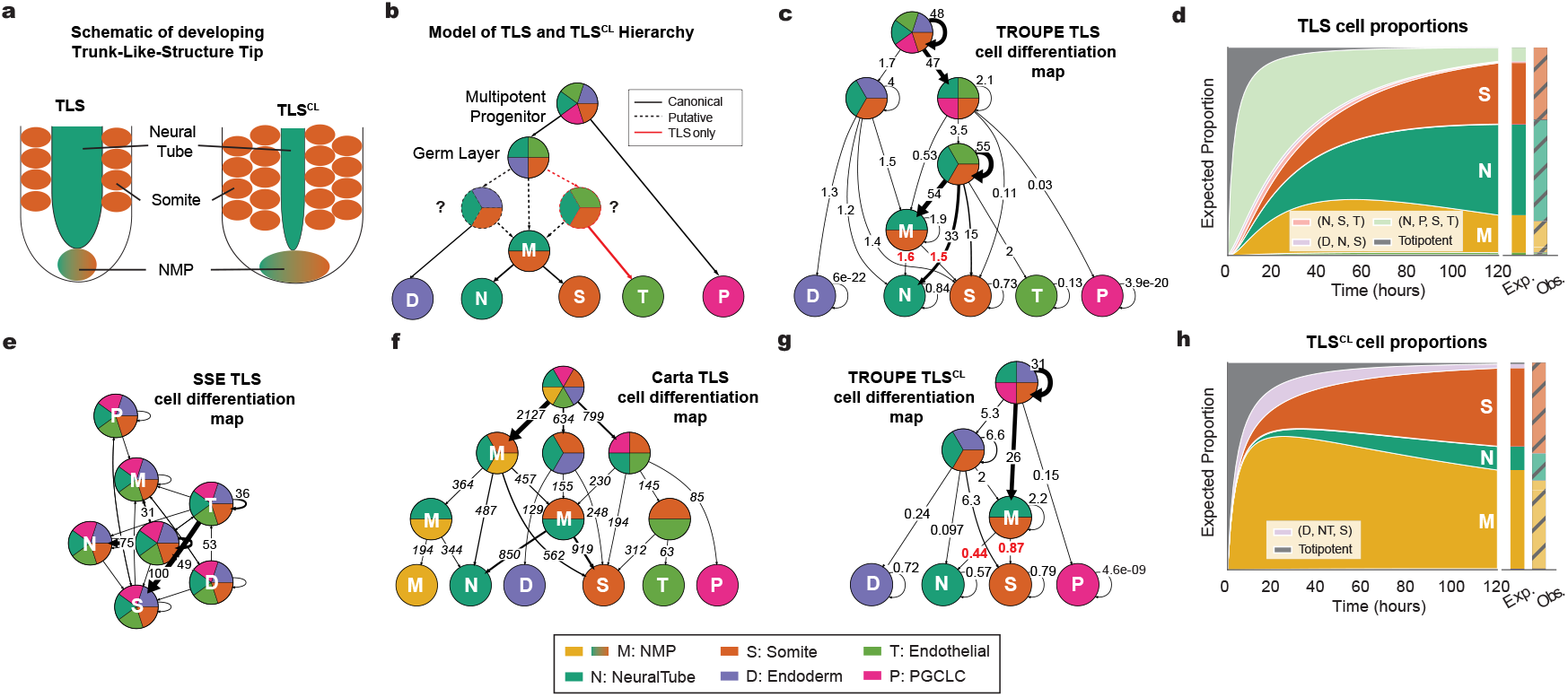
TROUPE infers dynamics of Trunk-Like Structures (TLS) model of mammalian development, highlighting differences between the standard condition as well as a chemically perturbed condition (TLS^CL^). (a) A schematic of the developing TLS trunk in the standard and TLS^CL^ conditions. The TLS^CL^ is expected to have an overgrowth of somites and a shrinking of the neural tube, as well as a larger pool of replicating NMPs. (b) The canonical cell differentiation map topology for TLS and TLS^CL^. The observed cell types include endoderm cells, somite cells, neuormesodermal progenitors (NMPs), neural tube cells, endothelial cells, and primordial germ-cell-like cells (PGCLCs). Dotted edges and nodes represent putative transitions and progenitors respectively. Red indicates progenitors and transitions only included in TLS but not TLS^CL^, as endothelials are not sampled in TLS^CL^. (c) The TROUPE TLS cell differentiation map with 5 progenitor cell types. Each node represents a row of ***Q***^*ω*^, the edge weight from node *u* ≠*v* is ***Q***_*u,v*_, and the self-edge weight for node *v* is given by ***ω***_*v*_. Red edge weights highlight the transition rates coming from NMP to neural tube and somite cell types. The unit of the rates are differentiation/division per length of experiment (120 hours) (d) The expected proportion of each cell type over the experimental timeframe, as calculated from the inferred rates in TLS. The first bar plot to the right of the 120 hour mark shows the expected proportions at 120 hours, the time of experimental observation. The second bar plot shows the true proportions of cell types observed in the data. (e) The SSE TLS cell differentiation map with 7 states. NMPs are treated as a terminal state in this method. (f) The *Carta* TLS cell differentiation map with 7 progenitors. Note that NMPs are treated as a terminal state in this method. Edge weights correspond to the number of extant cells in the cell lineage trees that undergo that transition. (g) The TROUPE cell differentiation map for TLS^CL^. (i) The expected proportion of each cell type over the experimental timeframe, as calculated from the inferred rates in i) TLS^CL^.

We applied TROUPE to 12 cell lineage trees generated by Bolondi et al. [18]. These cell lineage trees were reconstructed from lineage barcodes using LAML [15], which output ultrametric timed trees with branch lengths where the length of the tree is 1.0. Hence the rates can be interpreted as differentiation/division per length of experiment (120 hours). These cell lineage trees include 2521 extant sampled cells that are labeled by 6 observed cell types derived from the gene expression measurements: Endoderm (12 cells), Endothelial (25 cells), Primordial germ cell-like cells (PGCLCs; 5 cells), Somite (869 cells), Neural Tube (1216 cells), and NMP (378 cells). In this dataset, leaves of the cell lineage tree are labeled as NMPs due to self-renewing population of NMPs that exist when the structures are sampled at 120 hours. When running TROUPE, we use the explicit constraint that NMPs can become Somite and Neural Tube cells, encoding that these cells are a bipotent progenitor cell type. We applied TROUPE with a variety of regularization values (0, 0.001, 0.003, 0.01, 0.03, 0.1, 0.3, 1, 3, 10, 30) and produce the Pareto front as described in 3.4. This process gives a Pareto front with cell differentiation maps with 2, 3, 5, 7, 8 and 11 progenitors (Figure S2a). We determine that the optimal cell differentiation map has 5 progenitors.

The cell differentiation map inferred by TROUPE for the standard TLS model system agrees with known features of TLS development. The relative ordering of the commitment of cell types in the TROUPE cell differentiation map (Fig. 3c) is consistent with that of the canonical differentiation hierarchy of TLS (Fig. 3b). The PGCLCs differentiate closer to multipotency (the root) with an average progenitor potency size of 4.5. Endoderm and endothelial cells follow next in the differentiation hierarchy as they derive from progenitors with more restricted potencies (avg. size: 4.0), and the closely related somites and NMPs follow them (avg. size: 3.4). The TROUPE cell differentiation map suggests that there is indeed a separation of the multipotent pool as described above with the inclusion of both the {somite, endothelial, neural tube} and {somite, endoderm, neural tube} progenitors. The TROUPE and *Carta* (Figure 3f) cell differentiation maps have very similar topologies. Because the *Carta* framework does not encode observed cell types as progenitors, NMPs are also encoded via multiple progenitors ({ neural tube, NMP}, {somite, neural tube}, and {neural tube, somite, NMP}) in the *Carta* cell differentiation map. If these progenitors are collapsed and treated as one “NMP” progenitor, the only differing progenitors are the {somite, endothelial, neural tube} progenitor inferred by TROUPE and the {somite, endothelial} inferred by *Carta*. Both of these progenitors capture the shared origin of somite and endothelial cells [40, 41].

TROUPE provides an interpretable view of differentiation dynamics that other methods do not by inferring growth and transition rates. The dynamics of most interest in this system are the rates of NMP differentiation into somite and neural tube cells. TROUPE infers that the rate of differentiation of NMPs into either cell type is about equal (transition rate of 1.5 from NMPs to somites and 1.6 from NMPs to neural tube, Fig. 3c). In contrast, the dynamics of NMP differentiation are difficult to interpret from the cell differentiation map inferred by other methods. SSE with 7 states (one for each terminal state and one multipotent root state) does not accurately infer known features of the TLS system due to the lack of potency constraints and unobserved progenitors in the standard SSE model. An immediate incongruity is that it suggests terminal states that are not known to transition to each other have non-zero inferred transition rates (Fig. 3e). Hence, the rates inferred by SSE do not provide any cogent information about the differentiation dynamics in this system. As noted earlier, in the *Carta* cell differentiation map (Figure 3f), NMP dynamics are encoded through a complex series of possible progenitor types including the {neural tube, NMP}, {somite, NMP}, {somite, neural tube}, and {neural tube, somite, NMP} progenitors (Fig. 3f). It is difficult to interpret the difference between the progenitors in this hierarchy at different levels. Further, the edge weights in the *Carta* cell differentiation map represent the empirical proportion of cells that transition between each cell type and are not rates. Hence, they do not provide any information about the timing of these transitions and may not accurately reflect true probabilities of transition as they can be biased by the growth rates of subsequent cell types. *Carta* also does not give any characterization of self-renewal or growth as it does not include self-edges.

In addition to the cell lineage trees in the standard model of TLS development, we applied TROUPE to 11 cell lineage trees in the TLS^CL^ condition. These cell lineage trees consist of a total of 3165 extant sampled cells labeled in the same way as in the standard TLS condition: Endoderms (15 cells), Primordial germ cell-like cells (PGCLCs; 3 cells), Somites (1381 cells), Neural Tube cells (411 cells), and NMPs (1342 cells). Note that endothelial cells are not sampled in this condition. We applied in the same way as in the TLS condition TROUPE, using regularization values 0.003, 0.1, 0.3, 1, 3, 10 and 30 and choosing the cell differentiation map with 4 progenitors through the Pareto front (FigS2).

TROUPE has the unique capability discern subtle changes in dynamics caused by a perturbation to the system by comparing inferred rates in two conditions of a developmental system. Inferring the dynamics through rates in this case is especially important as the topology of the canonical cell differentiation map for TLS and TLS^CL^ are the same (with sampling of the endothelial cells). The shifts in the dynamics of NMP differentiation in the cell differentiation map inferred by TROUPE for the TLS^CL^ condition match the morphological expectation. First, Bolondi et al. [18] observed that NMPs are expected to maintain a larger self-renewing population in TLS^CL^ compared to TLS [18], as indicated by the increased growth (renewal) rate of NMPs in the TROUPE TLS^CL^ cell differentiation map (2.2) (Fig. 3g) compared to the growth rate of NMPs in the TROUPE TLS cell differentiation map (1.9) (Fig. 3c). Secondly, TROUPE infers that differentiation from NMPs to somites and neural tube cells is symmetric in the normal condition (with rates of 1.5 and 1.6 respectively) while it is asymmetric in the perturbed condition with a bias towards somites (rates of 0.87 and 0.44, respectively) (Fig. 3g). These trends can be further seen in the expected proportions of cell types as predicted by the rates of TLS and TLS^CL^ under the PoSSE model. Across the time-points of the experimental window (0 to 120 hours), there is a greater relative expected proportion of NMPs and somites and a lower relative expected proportion of neural tube cells in TLS^CL^ (Fig. 3h) than in TLS condition (Fig. 3d). As a further validation of these rates, the expected proportions of the cell types at 120 hours are close to those observed, except that some number of progenitors are expected to still linger at that time-point.

## 5 Discussion

TROUPE jointly estimates cell-type transition and growth rates for both observed and unobserved progenitor states from cell lineage trees under a novel branching-process model PoSSE. PoSSe encodes the irreversibility and potency constraints of developmental processes, which are missing from existing SSE models. We show that when transitions obey such constraints, transient progenitor types disappear over time, making it difficult to infer their rates or even their number from terminal measurements alone. In simulations, imposing these constraints improves the sample efficiency of rate inference compared to unconstrained approaches. We further introduce a model-selection procedure, inspired by Sashittal et al. [20], to infer the number of unobserved progenitors and find that it performs well across a range of simulated scenarios. More generally, our results indicate that constraining SSE rate matrices to reflect known structure can improve statistical efficiency, and similar ideas could be used to encode other inductive biases beyond potency.

There are two primary limitations of the model presented in this work and corresponding directions for future work. First, PoSSE assumes a time-homogeneous branching process, with transition and growth rates that are constant over time. However, systems of development are highly coordinated in time, and many features of development are time-inhomogeneous such as time-dependent fate bias of progenitor cells [18, 42] and time-varying rates of proliferation [43]. Extending PoSSE to allow for time-varying rates would make the model better aligned with known developmental biology. Second, we use a version of the SSE likelihood that assumes the observed tree is complete and thus does not accommodate subsampling or cell death. However, cell death is present in organismal development [44] and the observed leaves are often only a small fraction of the cell population at the time of measurement [18] (in the TLS data we estimate that 2–5% of extant cells are observed). One systematic effect of subsampling is that the growth rates of terminal types is underestimated because branch lengths of the leaves of a subsampled tree are longer than they would be without subsampling [45]. Previous work [27, 29] derives the likelihood in a way that accounts for cell-death and subsampling, but these formulations require numerical integration making optimization less stable and substantially slower for large numbers of cell-types. Integrating such likelihoods, or suitable approximations to them, into PoSSE would make the framework more realistic for developmental data sets with substantial cell death and incomplete sampling.

Despite these limitations, TROUPE shows that multi-replicate single-cell lineage tracing combined with transcriptional readout can be used to jointly infer both cell-type transition dynamics and type-specific growth rates under an explicit branching-process model. This demonstrates the value of adapting well-established models of population growth and state evolution to the context of differentiation at the single-cell resolution. We expect that TROUPE will be directly applicable to current lineage-tracing data sets and will provide a foundation for more detailed models and inference procedures as experimental technologies and developmental datasets continue to grow in scale and resolution.

## Code availability

Software is available at https://github.com/raphael-group/troupe

## 6 Acknowledgements

This work is supported by grant 2024-345885 from the Chan Zuckerberg Initiative DAF, an advised fund of the Silicon Valley Community Foundation; National Cancer Institute grant U24CA248453 to B.J.R.; National Institutes of Health grant DP2HD111537 to M.M.C; and the Princeton Catalysis Initiative. R.Z. is supported by National Human Genome Research Institute training grant T32HG003284.

## References

[1] Aaron McKenna, Gregory M Findlay, James A Gagnon, Marshall S Horwitz, Alexander F Schier, and Jay Shendure. Whole-organism lineage tracing by combinatorial and cumulative genome editing. Science, 353(6298): aaf7907, 2016.

[2] Young Seok Ju, Inigo Martincorena, Moritz Gerstung, Mia Petljak, Ludmil B. Alexandrov, Raheleh Rahbari, David C. Wedge, Helen R. Davies, Manasa Ramakrishna, Anthony Fullam, Sancha Martin, Christopher Alder, Nikita Patel, Steve Gamble, Sarah O’Meara, Dilip D. Giri, Torril Sauer, Sarah E. Pinder, Colin A. Purdie, Åke Borg, Henk Stunnenberg, Marc van de Vijver, Benita K.T. Tan, Carlos Caldas, Naoto Ueno, Laura J. van ‘t Veer, John W.M. Martens, Christos Sotiriou, Stian Knappskog, Paul N. Span, Sunil R. Lakhani, Jórunn Erla Eyfjörd, Anne L. Børresen-Dale, Andrea Richardson, Alastair M. Thompson, Alain Viari, Matthew E. Hurles, Serena Nik-Zainal, Peter J. Campbell, and Michael R. Stratton. Somatic mutations reveal asymmetric cellular dynamics in the early human embryo. Nature, 543(7647):714–718, 2017. doi: 10.1038/nature21703.

[3] Sara Bizzotto, Yanmei Dou, Javier Ganz, Ryan N. Doan, Minseok Kwon, Craig L. Bohrson, Sonia N. Kim, Taejeong Bae, Alexej Abyzov, the Brain Somatic Mosaicism Network, Peter J. Park, and Christopher A. Walsh. Landmarks of human embryonic development inscribed in somatic mutations. Science, 371(6535):1249–1253, 2021. doi: 10.1126/science.abe1544.

[4] Arikuni Uchimura, Hirotaka Matsumoto, Yasunari Satoh, Yohei Minakuchi, Sayaka Wakayama, Teruhiko Wakayama, Mayumi Higuchi, Masakazu Hashimoto, Ryutaro Fukumura, Atsushi Toyoda, Yoichi Gondo, and Takeshi Yagi. Early embryonic mutations reveal dynamics of somatic and germ cell lineages in mice. Genome Research, 32(5):945–955, 2022. doi: 10.1101/gr.276363.121.

[5] Michelle M. Chan, Zachary D. Smith, Stefanie Grosswendt, Helene Kretzmer, Thomas M. Norman, Britt Adamson, Marco Jost, Jeffrey J. Quinn, and et al. Molecular recording of mammalian embryogenesis. Nature, 570 (7759):77–82, 2019. doi: 10.1038/s41586-019-1184-5.

[6] Zhisong He, Ashley Maynard, Akanksha Jain, Tobias Gerber, Rebecca Petri, Hsiu-Chuan Lin, Malgorzata Santel, Kevin Ly, Jean-Samuel Dupré, Leila Sidow, et al. Lineage recording in human cerebral organoids. Nature methods, 19(1):90–99, 2022.

[7] Caleb Weinreb, Alejo Rodriguez-Fraticelli, Fernando D. Camargo, and Allon M. Klein. Lineage tracing on transcriptional landscapes links state to fate during differentiation. Science, 367(6479):eaaw3381, 2020. doi: 10.1126/science.aaw3381.

[8] Max A. Betjes, Sander J. Tans, and Jeroen S. van Zon. Cell tracking with accurate error prediction. bioRxiv, 2024. URL https://api.semanticscholar.org/CorpusID:273430743.

[9] Bushra Raj, James A Gagnon, and Alexander F Schier. Large-scale reconstruction of cell lineages using singlecell readout of transcriptomes and crispr–cas9 barcodes by scgestalt. Nature protocols, 13(11):2685–2713, 2018.

[10] Anna Alemany, Maria Florescu, Chloé S Baron, Josi Peterson-Maduro, and Alexander Van Oudenaarden. Wholeorganism clone tracing using single-cell sequencing. Nature, 556(7699):108–112, 2018.

[11] Matthew G. Jones, Azadeh Khodaverdian, Jeffrey J. Quinn, and et al. Inference of single-cell phylogenies from lineage tracing data using cassiopeia. Genome Biology, 21(92), 2020. doi: 10.1186/s13059-020-02000-8.

[12] Jean Feng, William S. Dewitt, Aaron Mckenna, Noah Simon, Amy D. Willis, and Fred-Erick A. Matsen. ESTIMATION OF CELL LINEAGE TREES BY MAXIMUM-LIKELIHOOD PHYLOGENETICS. The annals of applied statistics, 15(1):343–362, March 2021. ISSN 1932-6157. doi: 10.1214/20-aoas1400. URL https://www.ncbi.nlm.nih.gov/pmc/articles/PMC9387344/.

[13] Wuming Gong, Alejandro A. Granados, Jingyuan Hu, Matthew G. Jones, Ofir Raz, Irepan Salvador-Martínez, Hanrui Zhang, Ke-Huan K. Chow, Il-Youp Kwak, Renata Retkute, Alidivinas Prusokas, Augustinas Prusokas, Alex Khodaverdian, Richard Zhang, Suhas Rao, Robert Wang, Phil Rennert, Vangala Saipradeep, Naveen Sivadasan, Aditya Rao, Thomas Joseph, Rajgopal Srinivasan, Jiajie Peng, Lu Han, Xuequn Shang, Daniel J. Garry, Thomas Yu, Verena Chung, Mike J. Mason, Zhandong Liu, Yuanfang Guan, Nir Yosef, Jay A. Shendure, Maximilian J. Telford, Ehud Y. Shapiro, Michael B. Elowitz, and Pablo Meyer. Benchmarked approaches for reconstruction of in vitro cell lineages and in silico models of c. elegans and m. musculus developmental trees. Cell systems, 2021. URL https://api.semanticscholar.org/CorpusID:235486089.

[14] Palash Sashittal, Henri Schmidt, Michelle Chan, and Benjamin J. Raphael. Startle: A star homoplasy approach for CRISPR-Cas9 lineage tracing. Cell Systems, 14(12):1113–1121.e9, December 2023. ISSN 24054712. doi: 10.1016/j.cels.2023.11.005. URL https://linkinghub.elsevier.com/retrieve/pii/S2405471223003289.

[15] Gillian Chu, Uyen Mai, Henri Schmidt, and Benjamin J. Raphael. Maximum likelihood inference of time-scaled cell lineage trees with mixed-type missing data using LAML. Genome Biology, 26(1):189, July 2025. ISSN 1474-760X. doi: 10.1186/s13059-025-03649-9. URL https://doi.org/10.1186/s13059-025-03649-9.

[16] Weixiang Fang, Claire M. Bell, Abel Sapirstein, Soichiro Asami, Kathleen Leeper, Donald J. Zack, Hongkai Ji, and Reza Kalhor. Quantitative fate mapping: A general framework for analyzing progenitor state dynamics via retrospective lineage barcoding. Cell, 185(24):4604–4620.e32, November 2022. ISSN 00928674. doi: 10.1016/j.cell.2022.10.028. URL https://linkinghub.elsevier.com/retrieve/pii/S0092867422013745.

[17] Dian Yang, Matthew G. Jones, Santiago Naranjo, William M Rideout, Kyung Hoi Joseph Min, Raymond Ho, Wei Wu, Joseph M. Replogle, Jennifer L Page, Jeffrey J. Quinn, Felix Horns, Xiaojie Qiu, Michael Z. Chen, William A. Freed-Pastor, Christopher S. McGinnis, David M. Patterson, Zev J. Gartner, Eric D. Chow, Trever G. Bivona, Michelle M. Chan, Nir Yosef, Tyler Jacks, and Jonathan S. Weissman. Lineage tracing reveals the phylodynamics, plasticity, and paths of tumor evolution. Cell, 185:1905–1923.e25, 2022.

[18] Adriano Bolondi, Benjamin K Law, Helene Kretzmer, Seher Ipek Gassaloglu, René Buschow, Christina Riemen-schneider, Dian Yang, Maria Walther, Jesse V Veenvliet, Alexander Meissner, et al. Reconstructing axial progenitor field dynamics in mouse stem cell-derived embryoids. Developmental cell, 59(12):1489–1505, 2024.

[19] Kun Wang, Liangzhen Hou, Xin Wang, Xiangwei Zhai, Zhaolian Lu, Zhike Zi, Weiwei Zhai, Xionglei He, Christina Curtis, D. Zhou, et al. Phylovelo enhances transcriptomic velocity field mapping using monotonically expressed genes. Nature Biotechnology, 42(5):778–789, 2024.

[20] Palash Sashittal, Richard Y. Zhang, Benjamin K. Law, Alexander Strzalkowski, Henri Schmidt, Adriano Bolondi, Michelle M. Chan, and Benjamin J. Raphael. Inferring cell differentiation maps from lineage tracing data. bioRxiv, 2024. doi: 10.1101/2024.09.09.611835. URL https://www.biorxiv.org/content/early/2024/09/13/2024.09.09.611835.

[21] Joshua S. Schiffman, Andrew R. D’Avino, Tamara Prieto, Yakun Pang, Yilin Fan, Srinivas Rajagopalan, Catherine Potenski, Toshiro Hara, Mario L. Suva, Charles Gawad, and Dan A. Landau. Defining heritability, plasticity, and transition dynamics of cellular phenotypes in somatic evolution. Nature genetics, 56(10):2174–2184, October 2024. ISSN 1061-4036. doi: 10.1038/s41588-024-01920-6. URL https://www.ncbi.nlm.nih.gov/pmc/articles/PMC11527590/.

[22] Wayne P. Maddison, Peter E. Midford, and Sarah P. Otto. Estimating a Binary Character’s Effect on Speciation and Extinction. Systematic Biology, 56(5):701–710, October 2007. ISSN 1063-5157. doi: 10.1080/10635150701607033. URL https://doi.org/10.1080/10635150701607033.

[23] Richard G. FitzJohn. Diversitree: comparative phylogenetic analyses of diversification in R. Methods in Ecology and Evolution, 3(6):1084–1092, 2012. ISSN 2041-210X. doi: 10.1111/j.2041-210X.2012.00234.x. URL https://onlinelibrary.wiley.com/doi/abs/10.1111/j.2041-210X.2012.00234.x. eprint: https://besjournals.onlinelibrary.wiley.com/doi/pdf/10.1111/j.2041-210X.2012.00234.x. https://academic.oup.com/sysbio/

[24] Jeremy M. Beaulieu and Brian C. O’Meara. Detecting Hidden Diversification Shifts in Models of Trait-Dependent Speciation and Extinction. Systematic Biology, 65(4):583–601, July 2016. ISSN 1063-5157, 1076-836X. doi: 10.1093/sysbio/syw022. URL https://academic.oup.com/sysbio/article-lookup/doi/10.1093/sysbio/syw022.

[25] Richard G. FitzJohn. Quantitative Traits and Diversification. Systematic Biology, 59(6):619–633, December 2010. ISSN 1076-836X, 1063-5157. doi: 10.1093/sysbio/syq053. URL https://academic.oup.com/sysbio/article/59/6/619/1711291. https://academic.oup.com/sysbio/

[26] Richard G. FitzJohn, Wayne P. Maddison, and Sarah P. Otto. Estimating Trait-Dependent Speciation and Extinction Rates from Incompletely Resolved Phylogenies. Systematic Biology, 58(6):595–611, December 2009. ISSN 1076-836X, 1063-5157. doi: 10.1093/sysbio/syp067. URL https://academic.oup.com/sysbio/article/58/6/595/1635150.

[27] Denise Kühnert, Tanja Stadler, Timothy G. Vaughan, and Alexei J. Drummond. Phylodynamics with Migration: A Computational Framework to Quantify Population Structure from Genomic Data. Molecular Biology and Evolution, 33(8):2102–2116, April 2016. ISSN 0737-4038. doi: 10.1093/molbev/msw064. URL https://doi.org/10.1093/molbev/msw064. eprint: https://academic.oup.com/mbe/article-pdf/33/8/2102/7946826/msw064.pdf.

[28] Stilianos Louca and Michael Doebeli. Efficient comparative phylogenetics on large trees. Bioinformatics, 2017. doi: 10.1093/bioinformatics/btx701.

[29] Ailene MacPherson, Stilianos Louca, Angela McLaughlin, Jeffrey B Joy, and Matthew W Pennell. Unifying Phylogenetic Birth–Death Models in Epidemiology and Macroevolution. Systematic Biology, 71(1):172–189, June 2021. ISSN 1063-5157. doi: 10.1093/sysbio/syab049. URL https://www.ncbi.nlm.nih.gov/pmc/articles/PMC8972974/.

[30] T. Stadler, O. G. Pybus, and M. P. H. Stumpf. Phylodynamics for cell biologists. Science, 371(6526): eaah6266, January 2021. doi: 10.1126/science.aah6266. URL https://www.science.org/doi/full/10.1126/science.aah6266. Publisher: American Association for the Advancement of Science.

[31] Theodore E. Harris. The theory of branching processes. Die Grundlehren der Mathematischen Wissenschaften, Bd. 119. Springer-Verlag, Berlin, 1963.

[32] Krishna B. Athreya and Peter E. Ney. Branching Processes. Springer Berlin Heidelberg, Berlin, Heidelberg, 1972. ISBN 978-3-642-65373-5978-3-642-65371-1. doi: 10.1007/978-3-642-65371-1. URL http://link.springer.com/10.1007/978-3-642-65371-1.

[33] Roger S. Day. A branching-process model for heterogeneous cell populations. Mathematical Biosciences, 78(1): 73–90, March 1986. ISSN 00255564. doi: 10.1016/0025-5564(86)90032-5. URL https://linkinghub.elsevier.com/retrieve/pii/0025556486900325.

[34] Richard Durrett. Branching Process Models of Cancer. Springer, 2015. doi: 10.1007/978-3-319-16065-8.

[35] Einar Bjarki Gunnarsson, Jasmine Foo, and Kevin Leder. Statistical inference of the rates of cell proliferation and phenotypic switching in cancer. Journal of Theoretical Biology, 568:111497, July 2023. ISSN 00225193. doi: 10.1016/j.jtbi.2023.111497. URL http://arxiv.org/abs/2306.08096. arXiv:2306.08096 [q-bio].

[36] Dakota Dragomir, Elizabeth S. Allman, and John A. Rhodes. Parameter identifiability of a multitype pure-birth model of speciation. Journal of Computational Biology, 30(3):277–292, 2023. doi: 10.1089/cmb.2022.0330. URL https://doi.org/10.1089/cmb.2022.0330. PMID: 36745414.

[37] Dong C. Liu and Jorge Nocedal. On the limited memory bfgs method for large scale optimization. Mathematical Programming, 45:503–528, 1989. URL https://api.semanticscholar.org/CorpusID:5681609.

[38] Ville A. Satopaa, Jeannie R. Albrecht, David E. Irwin, and Barath Raghavan. Finding a “kneedle” in a haystack: Detecting knee points in system behavior. 2011 31st International Conference on Distributed Computing Systems Workshops, pages 166–171, 2011. URL https://api.semanticscholar.org/CorpusID:67623.

[39] Jesse V Veenvliet, Adriano Bolondi, Helene Kretzmer, Leah Haut, Manuela Scholze-Wittler, Dennis Schifferl, Frederic Koch, Léo Guignard, Abhishek Sampath Kumar, Milena Pustet, et al. Mouse embryonic stem cells self-organize into trunk-like structures with neural tube and somites. Science, 370(6522):eaba4937, 2020.

[40] Mounia Lagha, Silvia Brunelli, Graziella Messina, Ana Cumano, Tsutomu Kume, Frédéric Relaix, and Margaret E Buckingham. Pax3: Foxc2 reciprocal repression in the somite modulates muscular versus vascular cell fate choice in multipotent progenitors. Developmental cell, 17(6):892–899, 2009.

[41] Phong Dang Nguyen, Georgina Elizabeth Hollway, Carmen Sonntag, Lee Barry Miles, Thomas Edward Hall, Silke Berger, Kristine Joy Fernandez, David Baruch Gurevich, Nicholas James Cole, Sara Alaei, et al. Haematopoietic stem cell induction by somite-derived endothelial cells controlled by meox1. Nature, 512(7514): 314–318, 2014.

[42] Mina NF Morcos, Congxin Li, Clara M Munz, Alessandro Greco, Nicole Dressel, Susanne Reinhardt, Katrin Sameith, Andreas Dahl, Nils B Becker, Axel Roers, et al. Fate mapping of hematopoietic stem cells reveals two pathways of native thrombopoiesis. Nature communications, 13(1):4504, 2022.

[43] David R Kornack and Pasko Rakic. Changes in cell-cycle kinetics during the development and evolution of primate neocortex. Proceedings of the National Academy of Sciences, 95(3):1242–1246, 1998.

[44] M Suzanne and H Steller. Shaping organisms with apoptosis. Cell Death & Differentiation, 20(5):669–675, 2013.

[45] Tobias Dieselhorst and Johannes Berg. Branch length statistics in phylogenetic trees under constant-rate birth-death dynamics, October 2025. URL http://arxiv.org/abs/2407.13403. arXiv:2407.13403 [q-bio].

