## Supplementary Material for "Inferring Cell Differentiation Dynamics with Unobserved Progenitors"

### S6 Tree Construction

The descendant tree induced by a multi-type branching process can be viewed as sampling repeated independent *edge processes*. Each edge process starts with an initial type $s$ and samples a branch length $b$ and a coloring $c : [0, b] \to \Omega$ for that edge. In our setting, an edge process represents the cell-type transitions that a cell takes before dividing. Since each edge is sampled independently, describing the process for a single edge is sufficient for describing the process for all edges.

1. Initialize the starting time $\tau_0 = 0$ and the number of events that have occurred $k = 0$.
2. Let $s$ denote the current type. Sample time to next event $\tau_{k+1} \sim \exp\left(\lambda_s + \sum_{i \neq s} \boldsymbol{Q}_{si}\right)$.
3. Sample which event occurred $\mathbb{P}\,(\text{birth}) \propto \lambda_s$ and for $s \neq q$ we have $\mathbb{P}\,(s \text{ to } q) \propto \boldsymbol{Q}_{s,q}$.
4. Color the edge $c(t) = s$ for $\tau_k \leq t < \tau_{k+1}$ and increase the number of events that have occurred $k \leftarrow k + 1$.
    - If a $s$ transitions to $q$, replace the current type $s \leftarrow q$, and go back to step (2).
    - Otherwise, a birth occurs. Proceed to the next step.
5. Return the branch length $b = \sum_{i=1}^{k} \tau_i$ and the coloring $c$ for that branch.

Let $T = (V, E, b)$ be a length $t$ tree and let $\boldsymbol{y} : V \to \Omega$ be the state of the cell at each node. The tree $T$ and labeling $\boldsymbol{y}$ are generated as follows:

1. Initialize by adding root node $\rho$ to $V$, sampling the root state $\boldsymbol{y}_\rho \sim \boldsymbol{\pi}$, and setting the depth of the root to $d(\rho) = 0$.
2. For each current leaf node $u \in V$ such that $d(u) < t$, do the following steps once if $u$ is the root (to get the stem) and twice otherwise (i.e., once for each child):
    - Sample from edge process given starting type $\boldsymbol{y}_v$, which gives branch length $b$ and coloring $c$.
    - Add child node $v$ to $V$, edge $(u, v)$ to $E$.
    - If $b + d(u) > t$ (i.e., the edge would exceed the time limit), set $b = t - d(u)$ (i.e., truncate the branch).
    - Set $d(v) = d(u) + b$ and $\boldsymbol{y}_v = c(b)$.
3. Repeat (2) until all leaves have a distance to root of $t$.

This procedure produces a fully labeled, ultrametric tree of length $t$. In a lineage experiment, we only observe type information for the tips, so we let $\boldsymbol{x}_v = \boldsymbol{y}_v$ be the labels of just the leaf nodes $v \in V$. This tree-valued construction is equivalent to the one in Dragomir et al. [36], extended to include observed tip labels.

### S7 Proofs

#### S7.1 Vanishing Progenitors for two states

Proof of Proposition 2.1

*Proof.* For a two state branching processes, there is a simple closed form for the exponential of a matrix. Specifically,

$$\boldsymbol{M}(t) = \exp\left(t \cdot \begin{bmatrix} -\alpha + \lambda_1 & \alpha \\ 0 & \lambda_2 \end{bmatrix}\right) = \begin{bmatrix} e^{t(\lambda_1 - \alpha)} & \frac{e^{t\lambda_2} - e^{t(\lambda_1 - \alpha)})}{\alpha - \lambda_1 + \lambda_2} \\ 0 & e^{t\lambda_2} \end{bmatrix}. \quad (8)$$

Since $\alpha > \lambda_1$ by assumption, this formula holds for all valid $\alpha, \lambda_1, \lambda_2$. By linearity of expectation, the expected number of progenitor-type cells is $r\boldsymbol{M}(t)_{11}$. And we have

$$r\boldsymbol{M}(t)_{11} = r \exp\left(t(\lambda_1 - \alpha)\right)$$

by Equation (8). ☐

#### S7.2 Vanishing Progenitors for n states

Proof of Proposition 2.2.

We start by proving the following Lemma:

**Lemma S7.1.** *Let $M \in \mathbb{R}^{m\times m}$ be an upper triangular matrix with strictly negative diagonal entries. Additionally, assume that each of the diagonal entries of $M$ is unique. Then, $e^{tM} \to \mathbf{0}_{m\times m}$ as $t \to 0$.*

*Proof.* Observe that if $\boldsymbol{v} \in \mathbb{R}^m$ is an eigenvector of $M$ with eigenvalue $\gamma$, then

$$e^M \boldsymbol{v} = \left(\sum_{k=0}^{\infty} \frac{1}{k!} M^k\right) \boldsymbol{v} = \sum_{k=0}^{\infty} \frac{1}{k!} \gamma^k \boldsymbol{v} = e^{\gamma} \boldsymbol{v}$$

This implies that every eigenvector of $M$ with eigenvalue $\gamma$ is also an eigenvector of $e^M$ with eigenvalue $e^{\gamma}$. Moreover, for any upper triangular matrix $M$, so the eigenvalues of $M$ are exactly its diagonal entries $M_{ii}$. Combining this with the previous observation, we can see that the eigenvalues of $e^{tM}$ are $e^{tM_{ii}}$. Since $M_{ii} < 0$ we have that for the $i$th eigenvalue $\gamma = e^{tM_{ii}} \to 0$ as $t \to 0$. Since $M$ has $m$ unique, non-zero eigenvalues (i.e., its diagonal entries), it is diagonalizable. Let $VDV^\top = M$ be its diagonalization, where $D = \mathrm{diag}(M_{11}, ..., M_{mm})$. This gives us

$$\|e^{tM}\| = \|Ve^{tD}V^\top\| \le e^{t \max_i M_{ii}} \|V\|^2$$

Clearly, the right-hand side goes to 0 as $t \to \infty$ because $M_{ii} < 0$. This implies that $\lim_{t\to\infty} e^{tM} = \mathbf{0}_{m\times m}$. □

To see how this connects to expected population sizes of progenitor states in our branching process model, let $\boldsymbol{Q}'$, $\boldsymbol{\Lambda}'$ be the sub-matrices of $\boldsymbol{Q}$ and $\boldsymbol{\Lambda}$ that correspond to the progenitors in the process (i.e., rows of $\boldsymbol{Q}$ with non-zero diagonal entry). Without loss of generality, we can assume that these progenitors correspond to the first $m$ states in the rate matrix. That is,

$$\boldsymbol{Q} - \boldsymbol{\Lambda} = \begin{bmatrix} \boldsymbol{Q}' - \boldsymbol{\Lambda}' & \boldsymbol{A} \\ \mathbf{0} & \boldsymbol{B} \end{bmatrix}$$

for some matrices $\boldsymbol{A}, \boldsymbol{B}$. Exponentiating both sides gives

$$\exp(\boldsymbol{Q} - \boldsymbol{\Lambda}) = \begin{bmatrix} e^{\boldsymbol{Q}' - \boldsymbol{\Lambda}'} & \boldsymbol{C} \\ \mathbf{0} & e^{\boldsymbol{B}} \end{bmatrix}$$

for some matrix $\boldsymbol{C}$. Thus, the expected number of progenitors (given that we start from a progenitor type) is given by the rows of $e^{t(\boldsymbol{Q}'-\boldsymbol{\Lambda}')}$.

Note that $(\boldsymbol{Q}')_{ii} = -\sum_{j\neq i}^{n} \boldsymbol{Q}_{i,j} \le -\sum_{j\neq i}^{m} \boldsymbol{Q}_{i,j}$. Without loss of generality, we can assume that $\boldsymbol{Q}'$ is upper-triangular because the potency constraints enforce a DAG-like ordering on the rows. Therefore, $\boldsymbol{Q}' - \boldsymbol{\Lambda}'$ has negative entries on its diagonals. If we assume that the diagonal entries of $\boldsymbol{Q}' - \boldsymbol{\Lambda}'$ are unique, then the population of progenitor-types goes to 0 in the long time limit by Lemma S7.1.

#### S7.3 Potency-constrained MLE is equivalent to column selection

Proof of Proposition 3.1.

*Proof.* Let $\boldsymbol{Q}, \boldsymbol{\lambda}, P$ be a solution to Equation (5). Let

$$\boldsymbol{Q}' = \begin{bmatrix} \boldsymbol{Q} & \mathbf{0} \\ \mathbf{0} & \mathbf{0} \end{bmatrix} \quad \boldsymbol{\lambda}' = \begin{bmatrix} \boldsymbol{\lambda} \\ \mathbf{0} \end{bmatrix} \quad \boldsymbol{\Lambda}' = \mathrm{diag}(\boldsymbol{\lambda}')$$

Let $\Omega$ denote the labels of types that $\boldsymbol{Q}$ governs the transitions between and let $\Omega' \supseteq \Omega$ denote the set of labels that $\boldsymbol{Q}'$ governs the transitions between. Note that $\boldsymbol{Q}'$ is feasible to Equation (6) because it contains at most $n$ non-zero columns and inherits the potency constraints with respect to $P \subseteq \mathcal{P}(S)$.

Let $T = (V, E, b)$ be a tree with leaves labeled $\boldsymbol{x}$ and assume $\boldsymbol{x}$ contains more than two different labels. Let $\mathcal{Y}$ be the set of labelings of $V$ that extend $\boldsymbol{x}$. Let $\mathcal{Y}_0 \subseteq \mathcal{Y}$ be the set of labelings that only use labels corresponding to the first $n$

rows of $\boldsymbol{Q}$, and let $\mathcal{Y}_1 = \mathcal{Y} \setminus \mathcal{Y}$ the labelings that contain at least one label not from the first $n$ rows. For a full labeling $\boldsymbol{y} \in \mathcal{Y}$ of all the vertices (not just the leaves), let

$$L(T, \boldsymbol{y} \mid \boldsymbol{Q}, \boldsymbol{\lambda}) = \boldsymbol{\pi}_{y_\rho} \prod_{(u,v)\in E} \exp(b_{u,v}(\boldsymbol{Q} - \boldsymbol{\Lambda}))_{\boldsymbol{y}_u, \boldsymbol{y}_v} \prod_{v \in V_I} \lambda_{\boldsymbol{y}_v},$$

where $\boldsymbol{\pi}$ is fixed. We can partition the likelihood as

$$L(T, \boldsymbol{x} \mid \boldsymbol{Q}', \boldsymbol{\lambda}') = \sum_{\boldsymbol{y}\in\mathcal{Y}_0} L(T, \boldsymbol{y} \mid \boldsymbol{Q}', \boldsymbol{\lambda}') + \sum_{\boldsymbol{y}\in\mathcal{Y}_1} L(T, \boldsymbol{y} \mid \boldsymbol{Q}', \boldsymbol{\lambda}').$$

The second sum in this equation is 0. This holds because for any $t > 0$, we have

$$\exp(t(\boldsymbol{Q}' - \boldsymbol{\Lambda}')) = \begin{bmatrix} e^{t(\boldsymbol{Q}-\boldsymbol{\Lambda})} & \mathbf{0} \\ \mathbf{0} & \mathbf{I} \end{bmatrix} \tag{9}$$

Since the leaves are labeled by two different types, there must be an edge $(u, v) \in E$ such that $\boldsymbol{y}_u \in \Omega' \setminus \Omega$ and $\boldsymbol{y}_u \neq \boldsymbol{y}_v$. Any such transition has zero probability of occurring, therefore $L(T, \boldsymbol{y} \mid \boldsymbol{Q}', \boldsymbol{\lambda}') = 0$ for any $\boldsymbol{y} \in \mathcal{Y}_1$.

As for the first sum, any $\boldsymbol{y} \in \mathcal{Y}_0$ is such that

$$L(T, \boldsymbol{y} \mid \boldsymbol{Q}', \boldsymbol{\lambda}') = L(T, \boldsymbol{y} \mid \boldsymbol{Q}, \boldsymbol{\lambda}).$$

This is because $\boldsymbol{y}_v \in \Omega$ and thus, the transition probabilities are given by the upper-left block of the exponentiated matrix in Equation (9). Therefore,

$$L(T, \boldsymbol{x} \mid \boldsymbol{Q}', \boldsymbol{\lambda}') = L(T, \boldsymbol{x} \mid \boldsymbol{Q}, \boldsymbol{\lambda})$$

Let $\boldsymbol{Q}', \boldsymbol{\lambda}'$ be a solution to Equation (6). Let $k \leq n$ be the number of non-zero rows in $\boldsymbol{Q}'$. If column $i$ is zero, that implies that row $i$ is zero as well by the fact that we constrain $\boldsymbol{Q}'_{ii} = -\sum_{j\neq i} \boldsymbol{Q}_{ij}$. Without loss of generality, we can assume that

$$\boldsymbol{Q}' = \begin{bmatrix} \boldsymbol{Q} & \mathbf{0} \\ \mathbf{0} & \mathbf{0} \end{bmatrix}$$

for a potency constrained $k \times k$ rate matrix $\boldsymbol{Q}$. Let $\boldsymbol{\lambda}$ be the first $n$ entries of $\boldsymbol{\lambda}'$ (i.e., the ones that correspond to the types that $\boldsymbol{Q}$ transitions between). It suffices to show that $L(T, \boldsymbol{x} \mid \boldsymbol{Q}', \boldsymbol{\lambda}') = L(T, \boldsymbol{x} \mid \boldsymbol{Q}, \boldsymbol{\lambda})$, which can be done in a manner identical to before. □

### S8 TROUPE Algorithm

**Algorithm 1 Tree-informed Rate Optimization Under Potency Enforcement (TROUPE).**

1. For $\gamma_1 > \gamma_2 > \ldots > \gamma_k$
   (i) Solve Equation (7) to obtain $\hat{\boldsymbol{Q}}^{(\gamma)}, \hat{\boldsymbol{\lambda}}^{(\gamma)}$ and the induced potency set $P^{(\gamma)}$ consisting of the potencies of types reachable from the totipotent state.
   (ii) Debias by fixing $P^{(\gamma)}$ and refitting the potency-constrained MLE (Equation (5) with $P = P^{(\gamma)}$) to obtain $\tilde{\boldsymbol{Q}}^{(\gamma)}, \tilde{\boldsymbol{\lambda}}^{(\gamma)}$.
   (iii) Record model size $n^{(\gamma)}$ and fit $\log L\left(\mathcal{T} \mid \boldsymbol{Q}^{(\gamma)}, \boldsymbol{\lambda}^{(\gamma)}\right)$.
2. Form the pareto front of points among $\mathcal{C} = \left\{\left(n^{(\gamma)}, -L^{(\gamma)}\right) : \gamma \in \Gamma\right\}$.
3. Use Kneedle to identify the point $n^{(\gamma^\star)}$ of maximum curvature and return debiased estimates $\hat{\boldsymbol{Q}}^{(\gamma^\star)}, \hat{\boldsymbol{\lambda}}^{(\gamma^\star)}$.

### S9 Supplementary Figures

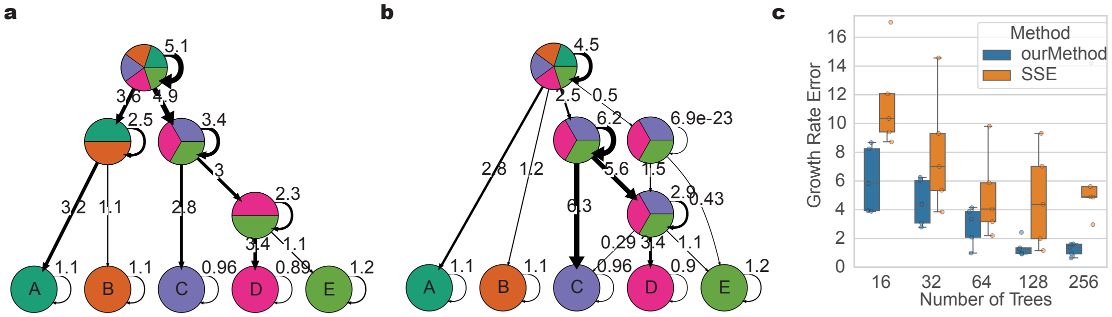


***Fig. S1*** **Additional plots regarding simulations.** a) An example cell differentiation map inferred by TROUPE for one trial in our simulations (32 trees). Each node represents a row of $\boldsymbol{Q}^{\star}$, the edge weight from node $u \neq v$ is $\boldsymbol{Q}_{u,v}$, and the self-edge weight for node $v$ is given by $\boldsymbol{\lambda}_v$. b) An example cell differentiation map inferred by SSE for one trial in our simulations (32 trees). c) Absolute error in growth parameter values for SSE and TROUPE for 5 trials for each number of trees.

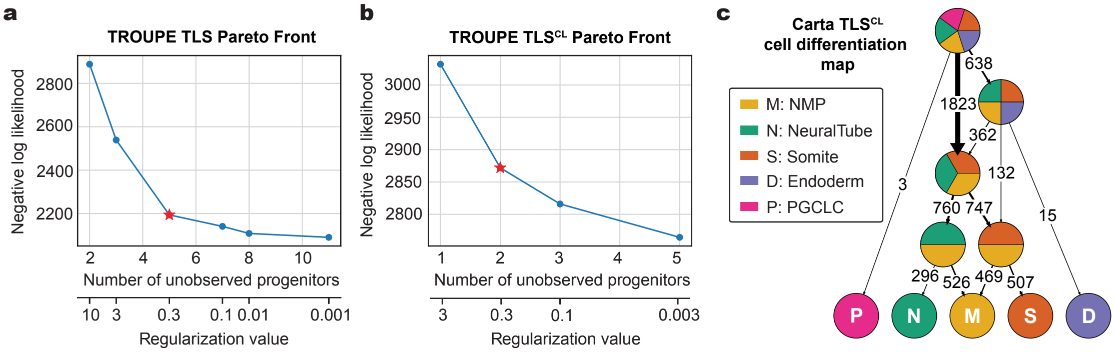


***Fig. S2*** **Additional plots regarding the Trunk-like Structures dataset.** The Pareto front showing the tradeoff between the negative log-likelihood and the number of states under various regularization values (0.001, 0.01, 0.1, 0.3, 3, 10) for a) TLS and b) $\text{TLS}^{\text{CL}}$. Star indicates the elbow point of the Pareto front, determined by Kneedle [38]. c) The *Carta* cell differentiation map with 5 progenitors. Note that NMPs are treated as a terminal state and endothelial cells are not present. Edge weights correspond to the number of extant cells in the cell lineage trees that undergo that transition.
